# Neat1 alleviates ischemic injury in skeletal muscles

**DOI:** 10.1101/2025.11.08.687205

**Authors:** Kyungmin Kim, Tao Yu, Hyejin Mun, Dong Gun Oh, Sangyoon Lee, Mirae Kim, Jibeen Kim, Yuseon Kang, Joshua Li, Young C. Jang, Je-Hyun Yoon, Laura Hansen, Hyojung J. Choo

## Abstract

Peripheral artery disease (PAD) has historically been regarded as an age-related vascular disorder; however, recent attention has shifted toward the myopathic components of the disease. Conventional interventions, such as revascularization, have had limited success in reversing muscle pathology or preventing adverse outcomes like amputation. Hypoxia-inducible factors (HIFs) are key regulators of cellular responses to ischemia, including the promotion of angiogenesis and glycolysis. While pharmacological stabilization of HIF proteins represents a promising therapeutic strategy, its efficacy is diminished in aged muscles due to their intrinsically low basal HIF expression.

We hypothesized that long noncoding RNAs (lncRNAs) might enhance the hypoxic response in aged muscle through post-transcriptional regulation of HIF expression. Our study identified a long noncoding RNA Neat1, as a critical mediator of the hypoxia-induced stress response, including upregulation of Hif1α. In a murine hindlimb ischemia model, Neat1 knockout mice exhibited extensive necrosis following femoral artery ligation, whereas Neat1 overexpression conferred protection against ischemic injury. Mechanistically, we found that Neat1 regulates the stability of *Hif1a* mRNA, as Hif1a transcript levels were significantly reduced in Neat1-deficient muscle cells. Importantly, aged muscles displayed a blunted hypoxic response due to diminished Hif1a expression—an effect that was reversed through Neat1 overexpression, resulting in improved resistance to ischemic damage.

In summary, our findings highlight Neat1 as a novel regulator of muscle adaptation to hypoxia in aging. Enhancing Neat1 expression may represent a promising therapeutic strategy for improving ischemic outcomes in patients with peripheral artery disease.

## Introduction

In the United States, there are more than 8.5 million people aged 40 and over with PAD, and 202 million people are affected globally^1^. Approximately 11% of these patients progress to chronic limb-threatening ischemia with pain at rest, gangrene/ulceration of tissue, and higher rates (∼20%) of limb amputation as well as mortality due to systemic multi-organ failure^2–4^. PAD patients suffer not only from insufficient blood flow to the tissue/organ but also from hypoxia-induced myopathies; however, current revascularization strategies do not prevent further complications sufficiently^5–9^. During limb ischemia and insufficient revascularization, the affected muscles cope with hypoxia by elevating levels of hypoxia-inducible factor 1/2 alpha (**HIF1/2α**), a master transcription factor necessary for angiogenesis and anaerobic glycolysis. While in normoxia, HIF proteins are rapidly degraded to half within 3 min by post-translational modification through a process called proline hydroxylation catalyzed by oxygen-sensing prolyl hydroxylases; during hypoxia, HIF1α protein is stabilized for up to 8 hours of half-life^10, 11^ due to lack of proline hydroxylation, and its activity on target gene transcription is sustained^12, 13^. Therefore, chemical inhibitors of HIF prolyl hydroxylases, also called HIF stabilizers, can boost hypoxic responses, such as angiogenesis and erythropoiesis. These chemicals such as Molidustat^14^, Vadadustat^15^, and Roxadustat^16^, are under clinical trials to treat anemia. In 2023, GSK1278863 (Daprodustat) has been approved for renal anemia treatment by the FDA^17^. These chemicals protect the heart and other organs during ischemia in preclinical studies^18, 19^. While the GSK1278863 protects muscles from contraction-induced injury, it fails to improve skeletal muscle function in PAD, due to insufficient increase of HIF proteins^20, 21^. Aged human and mouse muscles have lower levels of HIF1/2α and HIF1β^22, 23^, which implies poor hypoxic responses of aged muscles upon ischemic insults. In addition, the genetic loss of HIF proteins impairs muscle regeneration by dysregulation of satellite cells^23, 24^, muscle-specific stem cells essential for muscle regeneration. Given the high prevalence of PAD in the elderly, the decline of HIF1/2*α* and HIF1β levels in aged muscles must be addressed when developing new PAD therapies. Elevating HIF protein levels will be beneficial in combination with HIF protein stabilization strategies to ensure the hypoxic response is more effective in aged muscles.

Hypoxia globally changes transcriptional and post-transcriptional gene expression by accumulating proteins and RNAs in a discrete cellular compartment called RNA granules^25–29^. One of the RNA granules, nuclear paraspeckle, is a membrane-less organelle structured with lncRNA *nuclear paraspeckle assembly transcript 1* (*NEAT1*), which scaffolds core RNA-binding proteins^30–32^. Paraspeckles function in transcriptional regulation, RNA editing, and nuclear retention of RNAs and RNA-binding proteins^33, 34^. The roles of *Neat1* are well-recognized under stress conditions, such as cancer for its oncogenic roles and neurodegeneration in its protective roles^35, 36^. Relevant to hypoxia and glucose metabolism, Neat1 has been implicated in multiple cellular processes relevant to hypoxic adaptation. First, Neat1 regulates HIF1α expression, as genetic deletion in mice or knockdown in cultured cells reduces HIF1α mRNA and protein levels in cancer cells^37–39^. In addition, Neat1 promotes angiogenesis, enhancing the transcription of pro-angiogenic factors that support tumorigenesis^40, 41^. Neat1 also modulates glycolytic metabolism, where glucose deprivation and refeeding trigger cytosolic relocalization of the short Neat1_1 isoform, enabling it to form biochemical complexes with glycolytic enzymes PGK1, PGAM1, and ENO1^40^. Finally, Neat1 contributes to cell survival under hypoxia, as its overexpression protects both cancer and neural cells from hypoxia-induced apoptosis and enhances proliferation specifically under low-oxygen conditions^26, 42, 43^. Based on previous research and our strong preliminary data, we investigate the role of *Neat1* in PAD by using *in vivo* transgenic mice and aged mice with acute ischemic injury and *in vitro* cell models of primary myogenic progenitor cells with hypoxia.

## Materials and Methods

### Animals

C57BL/6J mice (strain #000664, 8–12 weeks old) were purchased from The Jackson Laboratory and housed under specific pathogen-free conditions. BALB/c mice (BALB/cAnNCrl, 10 weeks old, Charles River Laboratories) were used in select experiments. Aged mice (approximately 80 weeks old) were bred and aged in-house at Emory University. Neat1 knockout (KO) mice, originally generated by Dr. Shinichi Nakagawa at Riken Institute of Physical and Chemical Research^44^ were kindly transferred to Dr. Jiliang Zhao (August University) and Dr. Hyojung Choo (Emory University). Neat1 KO mice and wild-type littermates were used for experiments. All animal procedures were approved by the Institutional Animal Care and Use Committee (IACUC) at Emory University and were conducted in accordance with the guidelines of the National Institutes of Health.

### *In vivo* electroporation of skeletal muscle

*In vivo* electroporation was performed to deliver *Neat1_1* overexpression plasmid into the quadriceps muscle of the left hindlimb in both Balb/c and aged mice, based on a previously described protocol^45^. Mice were anesthetized with 2% isoflurane using a nose cone, and a single subcutaneous injection of sustained-release buprenorphine (0.1 mg/kg) was administered for analgesia. Fur covering the target quadriceps region was trimmed, and 40 μL of hyaluronidase solution (0.5 mg/mL, 10 units) was injected into the muscle using a Hamilton syringe to facilitate plasmid uptake by degrading extracellular matrix components. Two hours after hyaluronidase treatment, mice were re-anesthetized and 25 μL of *Neat1_1* overexpression plasmid (1–2 μg/μL in sterile PBS) or 25 μL of PBS for the control group was injected intramuscularly into the quadriceps using a Hamilton syringe with a 27G needle. Electroporation was immediately performed using an ECM830 electroporator (BTX, Harvard Apparatus) with the following parameters: low-voltage mode, 90 V, 20 ms pulse duration, 10 pulses, 480 ms interval, and unipolar polarity. Muscle contraction was visually confirmed with each pulse. Mice were allowed to recover on a heating pad after the procedure. Hindlimb ischemia surgery was performed on the limb one week after electroporation.

### Hindlimb ischemia (HLI) surgery

We used the procedure as we reported previously^46^. Mice were anesthetized using 2% isoflurane delivered via a nose cone. Buprenorphine (0.1 mg/kg, subcutaneous) was administered prior to the surgical procedure for analgesia. Under sterile conditions, a unilateral incision was made along the left thigh. The superficial femoral artery and vein were ligated with 6–0 silk sutures at two points: one proximal to the origin of the deep femoral artery, and the other just before the bifurcation of the tibial arteries. The intervening segment of the vessel was carefully removed, ensuring preservation of the femoral nerve. The incision site was then closed with a monofilament suture, and mice were placed on a warming pad during recovery.

### Necrosis scoring

Tissue necrosis was assessed daily and scored on a scale from 0 to 5 based on the extent of visible tissue damage and amputation as follows: 0 = no signs of necrosis; 1 = necrosis of a single toe; 2 = necrosis involving two or more toes; 3 = necrosis extending to the foot; 4 = necrosis of the leg and/or autoamputation of the foot; 5 = spontaneous loss of the entire limb (ref).

### Laser doppler perfusion imaging (LDPI)

Hindlimb perfusion was monitored noninvasively using a laser doppler perfusion imaging system (Moor Instruments) at the time points indicated in the figure following hindlimb ischemia (HLI) surgery. Mice were anesthetized with 2% isoflurane and placed on a heating pad to maintain body temperature. Hair on both hindlimbs was removed using depilatory cream and rinsed with water prior to imaging. After a 4-minute equilibration period, perfusion images were acquired. Consistent regions of interest were defined on both the ischemic hindlimb and the non-ischemic contralateral hindlimb, including whole, proximal, and distal segments. Perfusion was quantified in arbitrary perfusion units and expressed as the ratio of perfusion in the ischemic hindlimb to that in the non-ischemic hindlimb^46^.

### Aortic ring assay

Thoracic aortas were isolated from mice and cut into approximately 1-mm rings (ref). Each ring was embedded in a type I collagen matrix within a 96-well plate. Wells were treated with one of the following conditions: Opti-MEM alone (negative control), Opti-MEM supplemented with 0.3% vascular endothelial growth factor (VEGF, positive control), or conditioned media (Opti-MEM?) derived from wild-type or *Neat1* knockout muscle progenitor cells (MPCs) cultured under normoxic or hypoxic conditions. Conditioned media were collected after 24 hours of incubation using 2 x 10^6^ cells per T75 flask. Brightfield images were acquired using a Keyence microscope, and fluorescent images were captured with a confocal microscope. For quantitative analysis, total sprout length and the number of branching points were measured using ImageJ software. Brightfield and fluorescence images were analyzed independently.

### Muscle tissue preparation for histology

Quadriceps (Quad), tibialis anterior (TA), and gastrocnemius (GA) muscles were harvested from mice euthanized by an overdose of isoflurane. Freshly dissected muscles were embedded in Tissue-Tek® O.C.T. Compound (Sakura Finetek, Torrance, CA, USA), rapidly frozen in a bath of 2-methylbutane chilled with liquid nitrogen, and stored at −80 °C until sectioning. Transverse cryosections of 10 μm thickness were obtained using a CryoStar NX50 Cryostat (Thermo Fisher Scientific, Waltham, MA, USA) for subsequent histological analysis.

### Histological analysis

For general assessment of muscle morphology, cryosections were stained with hematoxylin and eosin (H&E) according to the manufacturer’s protocol. To examine fat infiltration, sections were immunostained with anti-perilipin A/B antibody (Sigma, P1873, 1:200) and FITC-conjugated wheat germ agglutinin (WGA, 1:50) to visualize muscle membranes. Sections were first incubated in blocking buffer containing 5% donkey serum, 0.5% bovine serum albumin (BSA), and 0.2% Tween-20 in PBS for 1 hour at room temperature, followed by overnight incubation at 4 °C with the perilipin A/B antibody. The next day, sections were washed three times with 0.2% Tween-20 in PBS and incubated for 1 hour at room temperature with Alexa Fluor® 594-conjugated donkey anti-rabbit IgG antibody (Jackson ImmunoResearch, 711-586-152, 1:200) along with FITC-WGA. To assess angiogenesis, sections were stained with anti-CD31 (PECAM1, BioLegend, 102405, 1:50) and anti-laminin antibody (Sigma, L9393, 1:200). After blocking as described above, sections were incubated with the laminin antibody for 1 hour at room temperature, followed by the same washing steps and subsequent incubation with Alexa Fluor® 594-conjugated donkey anti-rabbit IgG (Jackson ImmunoResearch, 711-586-152, 1:200) and FITC-conjugated CD31 antibody for 1 hour. Nuclei were counterstained with 4’,6-diamidino-2-phenylindole (DAPI, 1 μg/mL), and sections were mounted using VECTASHIELD mounting medium (Vector Laboratories). Images were acquired using an Echo Revolve widefield microscope, and quantitative analysis was performed using ImageJ software.

### Muscle Progenitor Cells (MPCs) isolation and culture

Muscle progenitor cells (MPCs) were isolated from hindlimb skeletal muscle of mice^47^. Dissected muscles were minced and enzymatically digested in 0.1% Pronase (EMD Millipore) diluted in Dulbecco’s Modified Eagle’s Medium (DMEM) supplemented with 25 mM HEPES buffer at 37 °C for 1 hour with gentle agitation at 150 rpm. Following digestion, the tissue was washed with DMEM containing 10% fetal bovine serum (FBS) and 100 μg/mL penicillin-streptomycin (Thermo Fisher Scientific). The cell suspension was then triturated using a 25 mL serological pipette and filtered through a 100 μm Steriflip filter (EMD Millipore) to obtain a mononuclear cell fraction. Cells were seeded on collagen-coated culture dishes and maintained in Ham’s F-10 medium (Thermo Fisher Scientific) supplemented with 20% FBS, 100 μg/mL penicillin-streptomycin, and 5 ng/mL basic fibroblast growth factor (bFGF; PeproTech). Cells were incubated at 37 °C in 5% CO₂ and atmospheric oxygen (∼20%) without media change for 3 days, after which MPCs morphology was confirmed. Once cells reached sufficient confluence, a pre-plating step was performed to enrich the MPCs population. Briefly, trypsinized cells were first plated on uncoated tissue culture dishes for 30 minutes to allow rapid attachment of non-MPCs populations. The supernatant containing non-adherent cells was collected and transferred to collagen-coated dishes for continued culture.

### Transfection

MPCs were transfected with plasmid DNA using Lipofectamine® 2000 reagent (Invitrogen, Thermo Fisher Scientific) according to the manufacturer’s instructions. Cells were seeded one day prior to transfection to reach 70–90% confluency at the time of transfection. For each well of a 6-well plate, 2 μg of plasmid DNA and 5 μl of Lipofectamine® 2000 were diluted separately in Opti-MEM® (Gibco) and incubated for 5 minutes before being combined and further incubated for 20 minutes at room temperature. The DNA–lipid complexes were then added to the cells in antibiotic-free medium and incubated for 4–6 hours, after which the medium was replaced with growth medium.

### Gene expression analysis by quantitative Real-Time PCR (qRT-PCR)

Total RNA was isolated from tissue or cell samples using TRIzol reagent (Invitrogen) according to the manufacturer’s protocol. For each sample, 200 ng of total RNA was reverse transcribed into complementary DNA (cDNA) using M-MLV reverse transcriptase (Invitrogen) with either random hexamers or oligo(dT) primers for *Neat1_1* amplification. Quantitative PCR was performed using Power SYBR® Green Master Mix (Applied Biosystems) and 2.5 μM of gene-specific primers on a QuantStudio 5 real-time PCR system (Applied Biosystems). Primer sequences are provided in Supplemental Table 1. Cycling conditions were as follows: 95°C for 15 seconds (denaturation), followed by 60°C for 1 minute (annealing/extension), for a total of 35 cycles. Expression of target genes was normalized to internal reference genes (*Gapdh* or *18S rRNA*), and relative gene expression levels were calculated using the comparative Ct method (ΔΔCt = (Ct^target^ - Ct^reference^)_sample - (Ct^target^ − Ct^reference^)_control) (Ref). Fold changes were derived using the equation 2^−ΔΔCt^. The PCR primers are listed in the Table S2.

### Immunoblotting

Cells were harvested and lysed in RIPA buffer (Thermo Fisher Scientific) supplemented with protease inhibitor tablets (Pierce, Thermo Fisher Scientific). Lysates were subjected to three cycles of freezing at −80 °C and thawing at room temperature, followed by vortexing and incubation on ice for 30 minutes. Insoluble material was removed by centrifugation at 21,000 × g for 15 minutes at 4 °C. Protein supernatants were mixed with Laemmli sample buffer (Bio-Rad) and boiled at 95 °C for 5 minutes. Samples were separated on 4–20% Mini-PROTEAN TGX Stain-Free gradient gels (Bio-Rad) and transferred to PVDF membranes. Membranes were blocked in 5% non-fat dry milk in TBS-T (Tris-buffered saline with 0.1% Tween-20) for 1 hour at room temperature and incubated overnight at 4 °C with primary antibodies diluted in 5% milk/TBS-T (antibodies listed in Supplementary Table S1). After three washes in TBS-T, membranes were incubated with HRP-conjugated secondary antibodies (1:5,000 dilution in 5% milk/TBS-T) for 1 hour at room temperature. Blots were developed using SuperSignal™ West Pico PLUS Chemiluminescent Substrate (Thermo Fisher Scientific) and visualized with a Kodak X-OMAT 2000A film processor.

### RNA pull-down using anti-sense DNA oligo

RNA pull-down using anti-sense DNA oligo was performed as follows. The biotinylated DNA ologos against *Hif1a* mRNA was designed and synthesized in IDT (Supplementary Table 2). MPCs lysates were prepared using protein extraction buffer containing 200 mM Tris-HCl pH 7.5, 100 mM KCl, 5mM MgCl2, 0.5% NP-40 (1 mg per sample), incubated with 1 μg of anti-sense or sense DNA oligo for 4 hours at 4°C. Complexes containing Hif1a mRNA were isolated with streptavidin-coupled Sepahrose beads (GE Healthcare, Chicago, IL, USA). The RNAs present in the pull-down material were analyzed by reverse transcription-quantitative polymerase chain reaction (RT-qPCR).

### Statistical Analyses

Statistical analysis was performed using Prism 10.0. Statistical testing was performed using the unpaired two-tailed Welch’s t-test to compare two groups or 1- or 2- way ANOVA depending on data structure, as stated in the figure legends. P-value <0.05 is considered statistical significance. Error bars represent the mean ± S.E.M.

## Results

### Neat1 deficiency aggravates muscle recovery after ischemic injury

To understand the roles of Neat1 in ischemic muscle injury, we performed femoral artery ligation in *Neat1^-/-^* and *Neat1^+/+^* (wild-type littermates) mouse muscles (3 months old in the background of C57BL/6, a strain well-tolerated ischemic injury after ligation^48^). While wild-type mice exhibited recovery of vessels, *Neat1^-/-^* mice failed to establish sufficient vessels (**Figure 1A-1B**) and developed tissue necrosis **(Figure 1C**). Muscle histology demonstrated significant changes in *Neat1^-/-^*. *Neat1^-/-^* mice have a smaller fiber size of the regenerated muscle by measuring cross-sectional area (CSA) of centrally nucleated myofibers but a higher fat infiltration by measuring the area of perilipin+ adipocytes in tibialis anterior muscles 28 days post-ischemic injury (**Figure 1D-1F**). Both indicate impaired muscle regeneration. In addition, *Neat1^-/-^*mice exhibited lower cells with PECAM-1^+^, a specific marker of angiogenesis 28 days post-ischemic injury (**Figure 1G-H**) in the tibialis anterior muscles. To investigate the molecular mechanism of poor recovery of *Neat1^-/-^* muscles from ischemic injury, we examined expressions of critical genes for coping ischemia, such as Hif1a, Hif2a, Vegfa, and Gapdh. We found that all those gene expressions were reduced in both injured and uninjured *Neat1^-/-^*gastrocnemius muscles, indicating that Neat1 may play a role in ischemic muscles by transcriptional regulation of those genes (**Figure 1I**). Taken together, Neat1 is critical for muscle recovery from ischemic injury.

**Figure 1.**
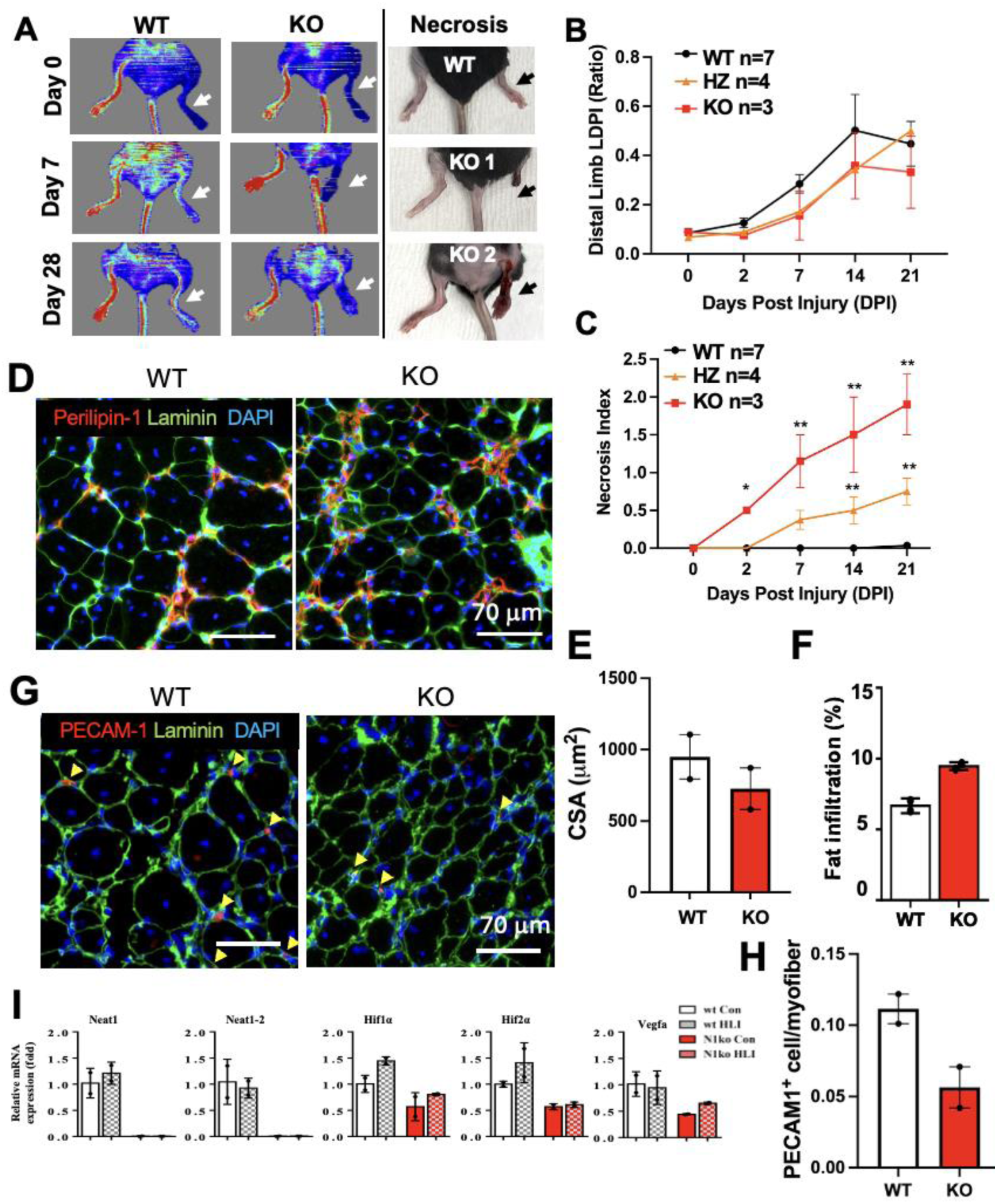
Neat1 deficiency exacerbates ischemic injury of limb muscles. **A.** Laser Doppler perfusion imaging (LDPI) of *Neat1^+/+^*(WT) and *Neat1^-/-^* (KO) mouse 0, 7 and 28 days following femoral artery ligation. Images of necrotic or loss of foot in *Neat1^-/-^*mouse post-ischemic injury. Arrows indicate injured hind limbs. **B-C.** LDPI*(B)* and Necrosis index *(C)* of *Neat1^+/+^, Neat1^+/-^* (HZ), and *Neat1^-/-^*mice by comparing ischemic distal limbs to control ones. **D.** Immunostaining of Perilipin-1 and Laminin with DAPI staining on tibialis anterior muscle sections 28 DPI (Days post injury). **E-F.** Quantification of (*A).* Cross-sectioned area (CSA) was measured by Laminin immunostaining, and fat-infiltrated areas were measured by Perilipin-1. n=2. **G.** Immunostaining of PECAM-1 and Laminin with DAPI staining on tibialis anterior muscle sections 28 DPI. **H.** Capillary number was counted by PECAM-1. n=2. **I.** Relative mRNA expression in gastrocnemius muscles of WT and *Neat1^-/-^* mice 28 days following femoral artery ligation.

### Muscle cells deficient in Neat1 fail to stimulate angiogenesis

Since Neat1^-/-^ muscles showed a delay in vessel recovery, we examined whether Neat1-mediated angiogenesis depends on the vessel- or muscle-derived factors. To test the angiogenic potential of *Neat1^+/+^* (WT) and *Neat1^-/-^* (KO) vessels, we isolated the aorta from adult mice and performed *ex vivo* mouse aortic ring angiogenesis assays with conditioned media (CM) of myogenic progenitor cells (MPCs) under hypoxia (**Figure 2A**). While we did not observe significant differences in the angiogenic potential of *Neat1^+/+^*and *Neat1^-/-^* aorta, CM of *Neat1^-/-^* MPC failed to stimulate the angiogenesis of *Neat1^+/+^* aorta (**Figure 2B-2C**). This indicates that impaired vessel recovery is derived from muscles or MPCs rather than the vessel itself.

**Fig. 2.**
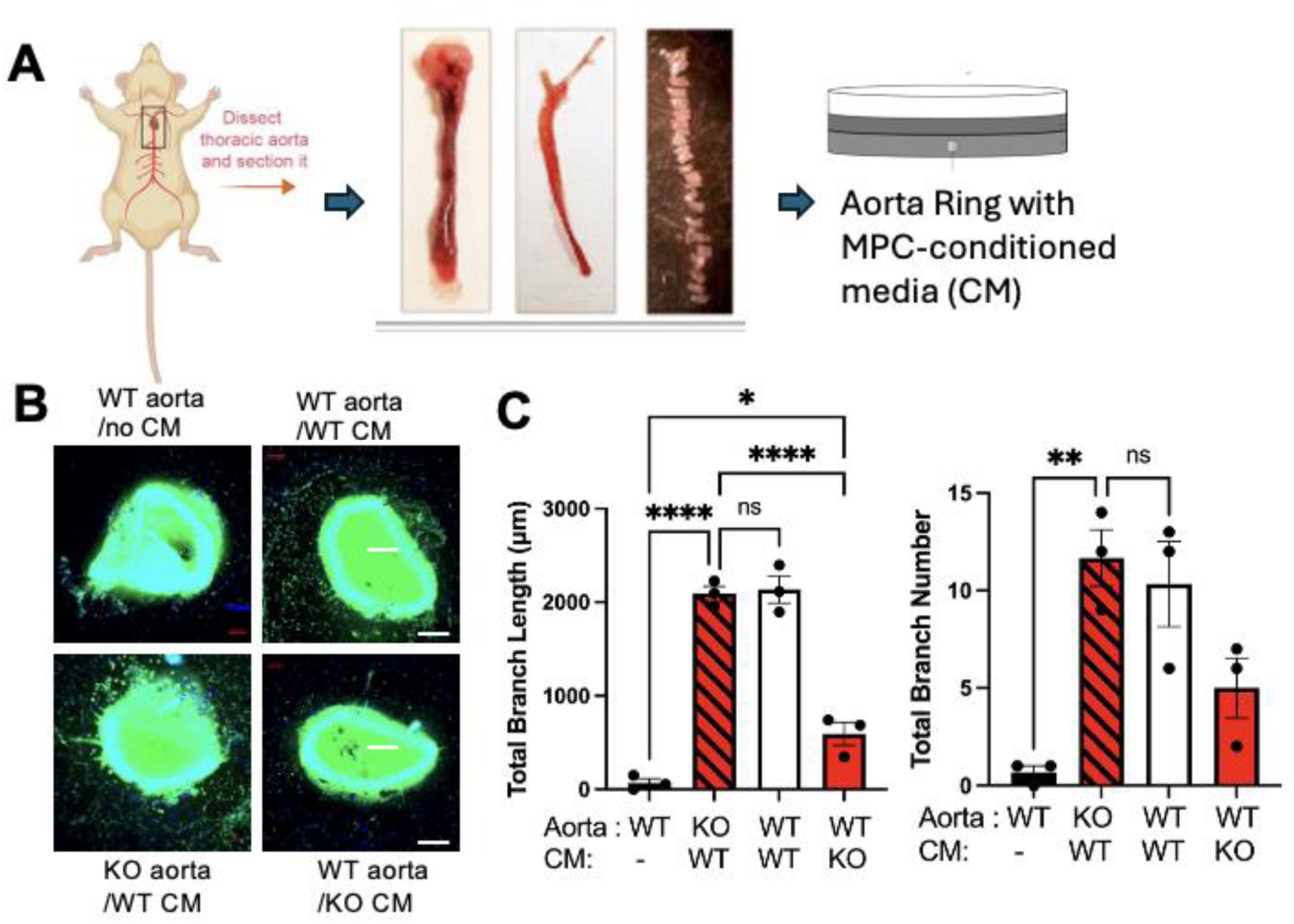
*Neat1* deficiency fails to stimulate MPC-derived angiogenesis. **A.** Experimental Scheme. **B.** Immunostaining images of the aorta from *Neat1^+/+^* (WT) and *Neat1^-/-^* (KO) mice with lectin to visualize sprouting and branching new vessels. Bar= 200μm. CM= conditioned media with WT or KO MPCs in hypoxia. **C.** Total branch length and number are measured and quantified to estimate angiogenesis of the aorta. n=3, Analyzed by 1-way ANOVA. **p<.01.

### Deficiency of Neat1_1 interferes hypoxic response of muscle cells

Our finding in *Neat1*-dependent changes of gene expression and muscle recovery post-ischemic injury, motivate us to investigate the molecular target of *Neat1*. During muscle hypoxia, the abundance of HIF1/2α mRNA and protein are elevated concomitantly. While both activate target gene transcription, such as *Vegfa* for angiogenesis, HIF1α mainly activates transcription of glycolytic genes, and HIF2α activates transcription of stem cell-related genes^49^. Moreover, HIF1α transcription is dependent on the presence of *Neat1* in cancers^37–39^.

Since muscles are composed of various cell types, our *in vivo* findings in ischemic injury and transcriptome changes could be driven by non-muscle cells surrounding muscle and secretory factors from gradual angiogenesis. To exclude those possibilities, we cultured *in vitro* primary myogenic progenitor cells (MPCs) and studied how Neat1 regulates the hypoxic response of muscles (1% O_2_, 48hr). As shown in **Figure 3A**, total *Neat1, Neat1_1, Neat1_2, Hif1α, Hif2α, Vegfα* (angiogenesis), and *Gapdh* (glycolysis) and as HIF downstream genes^50, 51^, were induced in wild-type MPCs upon hypoxia; however, those genes were not induced in *Neat1^-/-^* (both isoform knockout) MPCs. To distinguish the effect of individual *Neat1* isoform on the gene expression changes, we cultured myogenic progenitors isolated from *Neat1_1* (short Neat1 isoform with polyA tail)-knockout mice by deletion of poly-A site (**PAS**) on the Neat1 transcript, referred *dPAS-Neat1* allele. In *dPAS-Neat1* muscle cells, hypoxia-induced levels of *Hif1α, Hi2α, Vegfα,* and *Gapdh* were not observed anymore (**Figure 3B**). The most critical contributor to HIF activity in hypoxia is to escape HIF from degradation. Considering that HIF1α protein half-life is up to 8 hours^10, 11^, continuous transcription and/or attenuated mRNA decay would maintain HIF protein levels. Immunoblotting of Hif1α by using lysates from myogenic progenitor cells cultured under normoxia and hypoxia revealed elevation of HIF1α protein level in wild-type MPCs, but it significantly declined in *Neat1*-deficient cells under hypoxia (**Figure 3C**). Taken together, Neat1 deficiency in MPC causes lack of response to hypoxia via inhibiting Hif1a transcription.

**Fig. 3.**
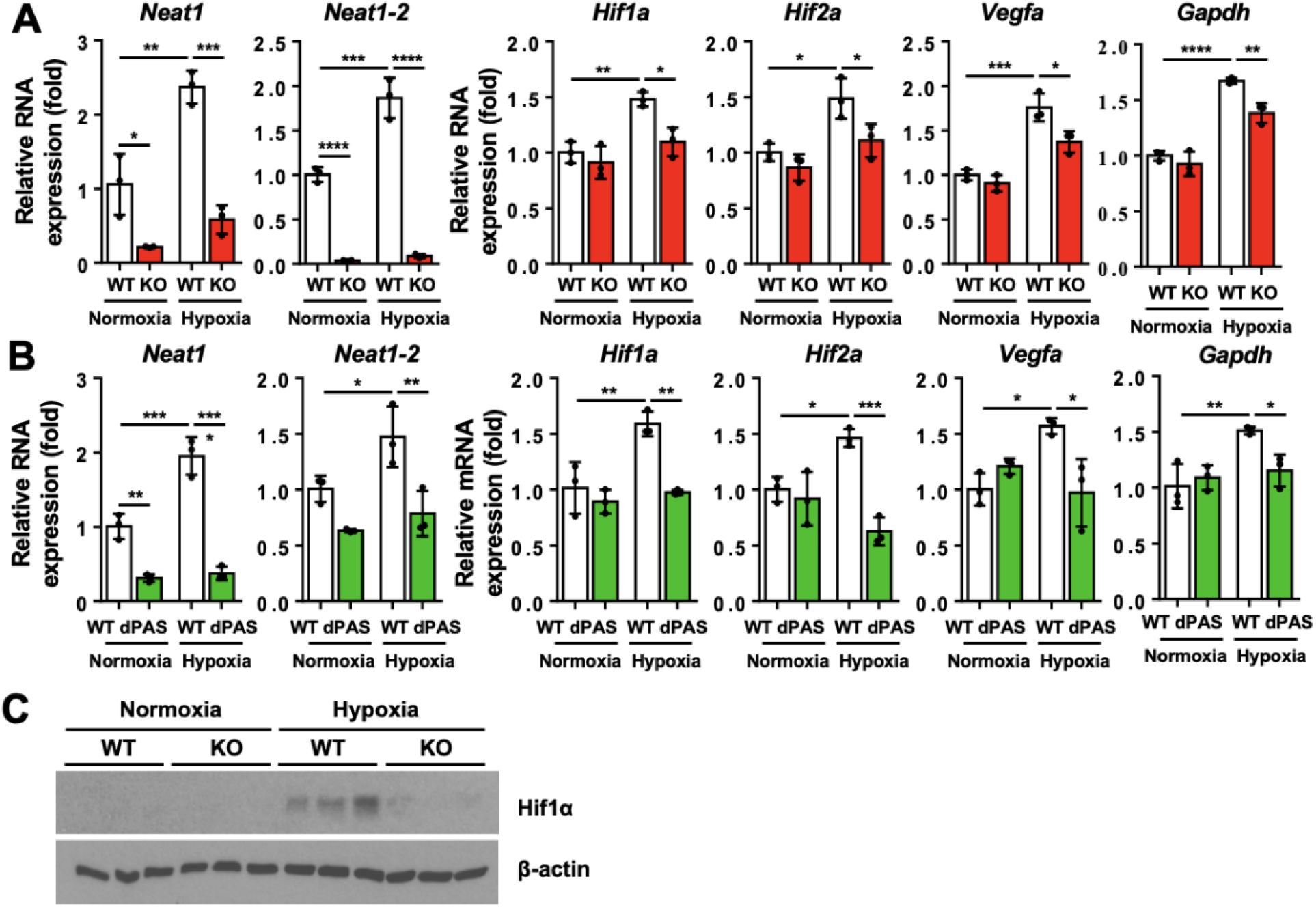
Neat1 is necessary to induce levels of *HIF* mRNA and protein in MPCs under hypoxia. **A-B.** RT-qPCR levels of total *Neat1, Neat1_2, Hif1a, Hif2a,* and *Vegfa* mRNAs in total RNAs isolated from cultured MPCs of *Neat1^+/+^* (WT) and *Neat1^-/-^* (KO) mice (*A*), *Neat1^+/+^* and *Neat1_1^-/-^* (dPAS) MPCs (*B*) under normoxia and hypoxia. **C.** Immunoblot levels of HIF1α and β-actin by using lysates of MPCs isolated from *Neat1^+/+^* and *Neat1^-/-^*under normoxia and hypoxia.

### Neat1_1 overexpression improves the hypoxic response of muscle cells

To investigate whether *Neat1_1* overexpression improves muscle recovery from ischemic injury, we used wild-type Balb/c mice, a strain that is susceptible to ischemic injury due to poor revascularization^48^, and overexpressed *Neat1_1* by in vivo electroporation on quadriceps muscles (***Neat1* Ox**). *Neat1_1* overexpression by electroporation before ischemic injury resulted in elevation of *Hif1α* mRNA level (**Figure 4A**), prompt vessel recovery, and less necrosis until day 7 (**Figure 4B-4D**), but the beneficial effect is reduced at day 14 after ligation, possibly due to decay of Neat1 plasmid (Data not shown). *Neat1_1* overexpression in wild-type Balb/c mice prevented muscle atrophy 7 days post-ischemic injury due to enhanced angiogenesis (**Figure 4E-4F**). Hif1a mRNA transcript level was increased by Neat1_1 overexpression (**Figure 4G)**

**Figure 4.**
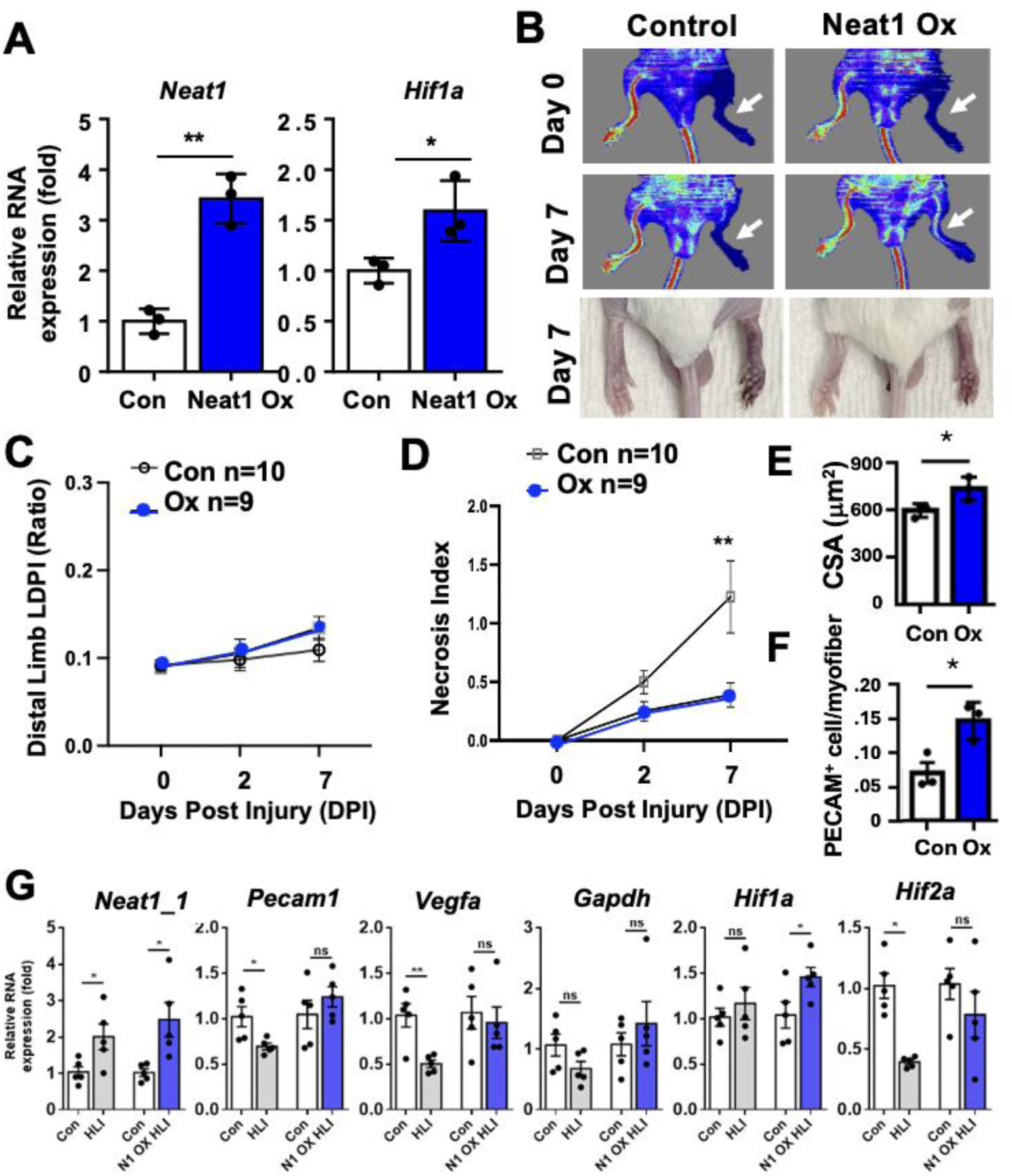
Neat1_1 overexpression improves recovery from ischemic injury of limb muscles. **A.** Relative RNA levels after *in vivo* electroporation of *Neat1_1* cDNA plasmid (*Neat1* Ox) or empty vector (Con) in quadriceps muscles (n=4). **B-D.** LDPI and foot pictures (*B*), and LDPI ratios (*C*), and necrosis index (*D*) of hind limbs 7 days following femoral artery ligation from wild-type Balb/c mice. **E.** Cross-sectioned area (CSA) was measured by Laminin immunostaining, and PECAM^+^ endothelial cells were counted by PECAM immunostaining. n=3. **F.** Relative mRNA expression in quadriceps muscles of WT and *Neat1_1-* overexpressed mice 7 days following femoral artery ligation. Analyzed by *t*-test or 2-way ANOVA. *p<.05, **p<.01

We transfected *Neat1_1* to MPCs to determine whether *Neat1_1* overexpression impact the expression levels of hypoxia responding genes (**Figure 5A**). *Neat1_1* overexpression induced the elevation of *Hif1α*, *Hif2α*, *Vegfα*, and *Gapdh* mRNAs upon hypoxia, which is opposite to Neat1-/- MPCs. These findings indicate that *Neat1_1* is a central player in regulating *Hif1/2α, Vegfα*, and *Gapdh* mRNA abundance. Also, HIF1α protein level is increased more in *Neat1_1*-overexpressed cells under hypoxia (**Figure 5B**). We also confirmed that human *HIF1A* is also regulated by *NEAT1* in **human MPCs** (**Fig. 3C**). Therefore, Neat1_1 is sufficient to induce Hif1/2α transcription, which supports sustained HIF proteins in skeletal muscles under hypoxia.

**Fig. 5.**
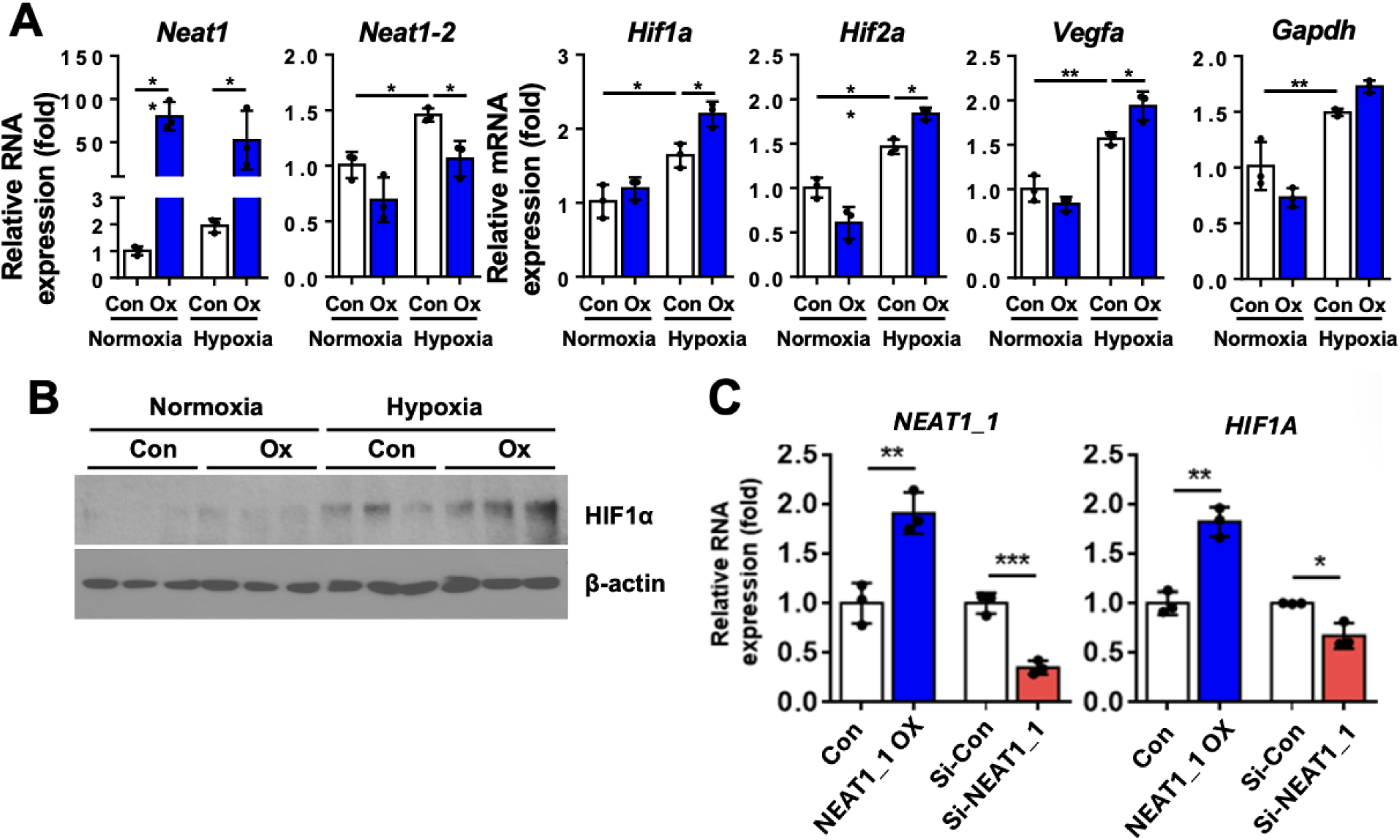
Neat1 is sufficient to induce levels of *HIF* mRNA and protein in MPCs under hypoxia. **A.** RT-qPCR levels of total *Neat1, Neat1_2, Hif1a, Hif2a,* and *Vegfa* mRNAs in total RNAs isolated from cultured MPCs of *Neat1^+/+^* (WT) and *Neat1_1* transfected (Ox) under normoxia and hypoxia. **B.** Immunoblot levels of HIF1α and β-actin by using lysates of MPCs with or without *Neat1_1* transfected under normoxia and hypoxia. **C.** RT-qPCR levels of *NEAT1_1 and HIF1A* from human MPCs transfected with *NEAT1_1 plasmid or si-NEAT1.* n=3, Analyzed by 2-way ANOVA. *p<.05, **p<.01.

### Neat1_1 overexpression improves the hypoxic response of aged muscles

Aging reduces HIF protein activity in muscles even under ischemia, making them susceptible to hypoxic stress and PAD-associated myopathies. Our data show that *Neat1_1* overexpression itself increases the abundance of *Hif1a* mRNAs in MPCs and *in vivo* muscle tissues, demonstrating its potential in enhancing the cellular response to ischemic injury and muscle regeneration of aged muscles via HIF regulation. To examine this possibility, we performed hind limb ischemic injury with *Neat1_1* overexpression by *in vivo* electroporation in old C57BL/6 mice, serving as a model for critical limb ischemia based on poor recovery in studies by our team (**Fig. 6A-C**) and others^52^. 18-month-old C57BL/6 mice with *Neat1_1* overexpression in the muscle had improved LDPI and reduced necrosis index 7 days post limb ischemic injury. (**Fig. 6D-F**). Interestingly, *Neat1_1* expression as well as *Hif1a* and HIF downstream gene expressions were reduced after 7 days post limb ischemic injury in aged muscles. *Neat1_1* overexpression recovered the impaired *Hif1a* and HIF downstream gene responses to ischemia in aged muscle. Particularly, angiogenesis marker gene (*Pecam1*) and myogenesis marker genes (eMyHC, *Pax7, MyoD, Myomaker)* are highly expressed (**Fig. 6G**).

**Fig. 6.**
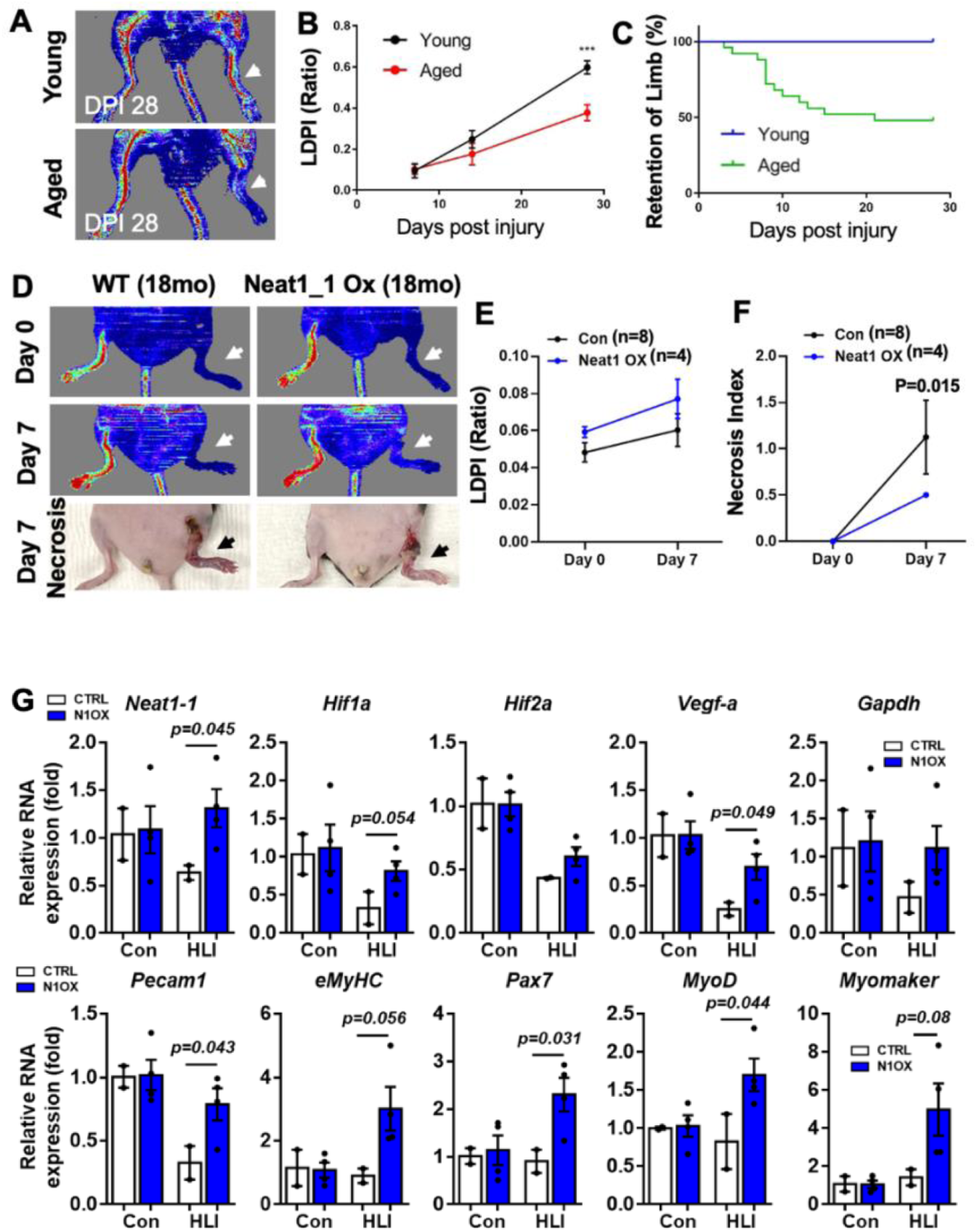
Neat1_1 overexpression reverses the impaired muscle recovery of aged muscles post-ischemic injury. **A-C.** LDPI outputs *(A)*, ratios *(B)* and Quantitation of limb retention (%) *(C)*. n=5. **D-F.** LDPI outputs and necrosis images and quantification of 18-month-old mice with femoral artery ligation after electroporation of *Neat1_1* plasmid or empty vector. n=4-8. **G.** RT-qPCR of total RNAs purified from lysates of quadriceps muscles from old mice with 7 days post-hind-limb ischemic injury (HLI) with or without *Neat1_1* overexpression. n=2-4. Analyzed by 2-way ANOVA. ***p<.001.

Similarly, we found that MPCs from old mice (24 months-old) failed to induce *Hif1a and Hif2a* mRNAs as well as *Neat1* upon hypoxia (**Fig. 7A; Nor**moxia Con vs **Hy**poxia Con), in contrast to MPCs from young mice (**Fig. 5A-B**). Treatment of HIF stabilizer^53^ (Roxadustat, 30μM, 3 hr.) in MPC from aged mice failed to accumulate HIF1a (**Fig. 7B**; Con), but *Neat1_1* overexpression made HIF stabilizer fully functional to induce Hif1a proteins in MPC from aged mice, even in normoxia conditions (**Fig. 7B**; N1OX). Taken together, our data indicate that Neat1-stabilized *Hif1a* mRNA aids strong HIF expression in aged muscles.

**Fig 7.**
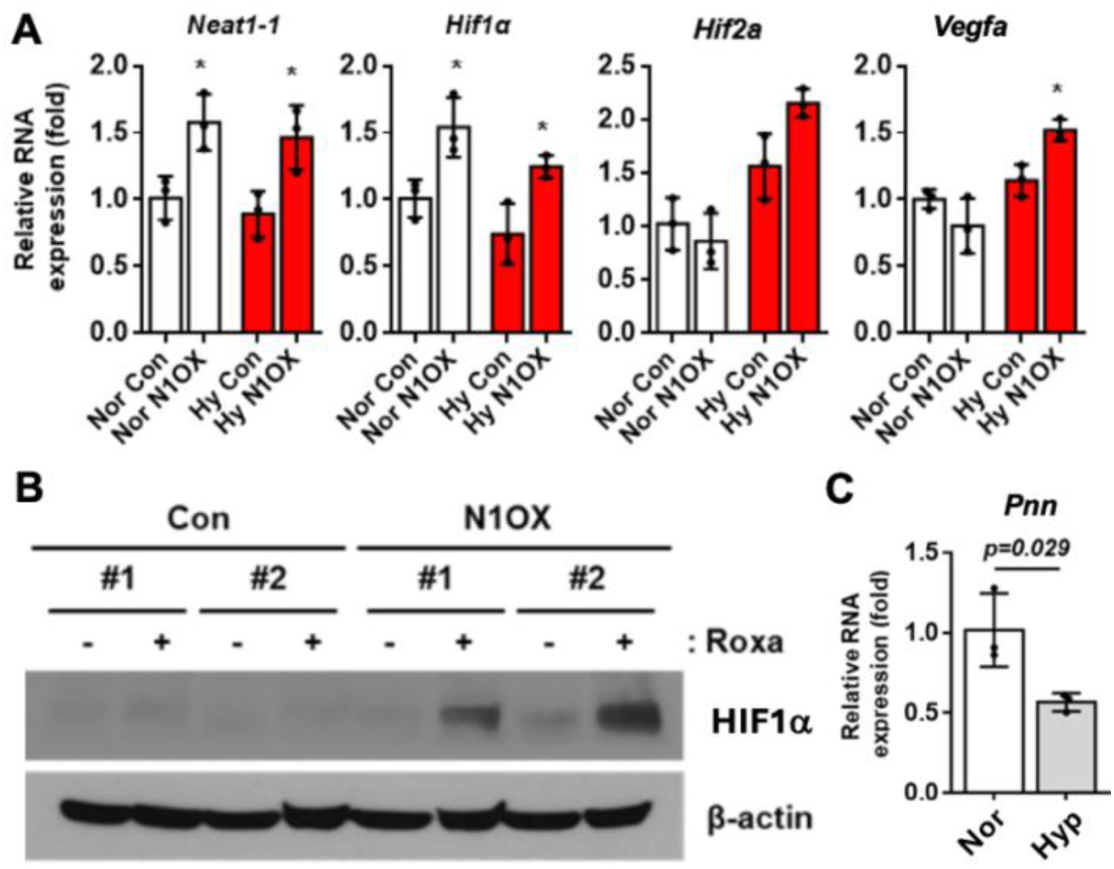
Impaired Hif1/2a induction upon hypoxia in aged MPCs. **A.** RT-qPCR levels of Neat1_1, *Hif1a*, *Hif2a*, and *Vegfa* mRNAs using MPCs lysates from old mice with normoxia or hypoxia (n=3). **B.** immunoblot levels of HIF1a and b-actin (loading control) with Roxadustat (30μ M, 3 hr.) in MPCs transfected with Neat1_1 cDNA plasmid. **C.** RT-qPCR levels of *Pnn* mRNA from samples used in *(A)* (n=3).

### Neat1 stabilizes *Hif1a* mRNA

To determine how Neat1 regulates Hif1a transcripts, we examine the possibility of direct binding between *Neat1* and *Hif1a* mRNA. To screen the transcripts binding with *Hif1a* mRNA, we computationally searched RNA sequences complementary to *Hif1a* mRNA from mouse transcriptome and found total 12 transcripts including 9 mRNAs and 3 lncRNAs (**Table 1**). To determine if *Hif1a* mRNA indeed binds those transcripts upon hypoxia, we pulled down *Hif1a* mRNA with its anti-sense oligonucleotides from myogenic progenitor cells (MPCs). As marked in **Table 1**, 2 mRNAs (Rcor3 and Vps37a) and 1 lncRNA (Neat1) showed higher enrichment with *Hif1a* mRNAs upon hypoxia (1% O_2_, 48 hr.). Since Neat1 contains the longest complementary sequences against *Hif1a* mRNA and protects mammalian cells from hypoxia, we prioritize Neat1 as the strongest candidate of the proposed studies.

**Table 1.** List of RNAs binding to Hif1a mRNA.

| lnc RNA | mRNA |
| --- | --- |
| Neat1*,<br>Gm31349,<br>Gm53955, | Mindy2, Fbxw10, Pabpc4,<br>Vldlr, **Rcor3\***, **Vps37a\***,<br>Arl5a, Boll, Cep20, Plet1os,<br>Dscam |

Among the isotypes of Neat1, Neat1_1 is specifically enriched with *Hif1a* mRNAs upon hypoxia (**Fig. 8A**), indicating a distinct role of each Neat1 isotype. To understand if Neat1 regulates *Hif1a* mRNA abundance and decay, we measured the half-life of *HIF* mRNA by shutting off transcription using an RNA polymerase inhibitor, Actinomycin D (**Fig. 8B**). Hypoxia stabilizes *Hif1a* mRNA in WT MPC but it was not observed in Neat1 KO MPC.

**Fig. 8.**
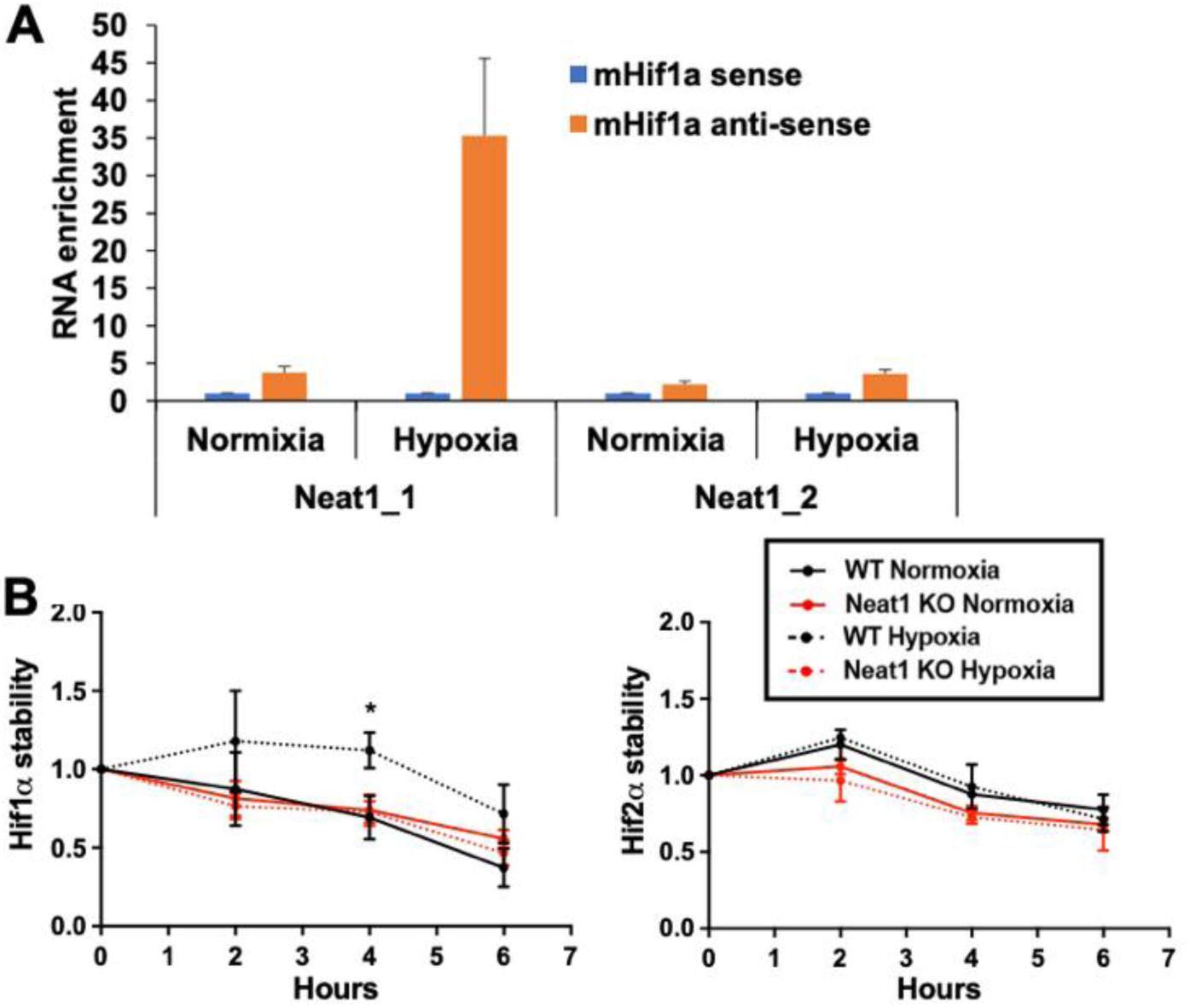
*Neat1* binds and stabilizes *Hif1a* mRNA. **A.** Affinity pull-down of *Hif1a* mRNA by using anti-sense or sense oligo enriches Neat1_1 in WT MPCs upon hypoxia (1% O_2_, 48hr) *in vitro*. **B.** RT-qPCR of *Hif1a and Hif2a* mRNAs of *Neat1^+/+^* (WT) and *Neat1^-/-^* (KO) MPC in normoxia and hypoxia (1% O_2_, 48hr) with Actinomycin D (2.5 μM).

## Discussions

Our study identifies Neat1, particularly the Neat1_1 isoform, as a key regulator of hypoxic adaptation and regenerative capacity in skeletal muscle during aging. We show that Neat1 directly binds to and stabilizes Hif1a mRNA, enabling sufficient induction of HIF1α protein and downstream transcriptional programs in muscle progenitor cells (MPCs) under hypoxic stress. Functionally, Neat1_1 overexpression enhances muscle recovery following ischemic injury, improves hypoxia-responsive gene expression, and promotes angiogenic signaling, restoring impaired responses observed in aged muscle. Conversely, loss of Neat1_1 disrupts hypoxic gene expression, reduces MPC-derived angiogenic capacity, and exacerbates ischemic muscle injury, demonstrating that Neat1_1 is required to maintain effective muscle regeneration and vascular remodeling in low-oxygen environments. Together, these findings position Neat1_1 as a crucial molecular mediator linking age-associated transcriptional decline to impaired ischemic adaptation in skeletal muscle.

Despite the significance of these results, several limitations should be considered. Our work primarily focuses on muscle progenitor cells, and the role of Neat1 in other muscle-resident cell types (e.g., endothelial cells, macrophages, fibro-adipogenic progenitors) during ischemic repair remains unclear. Additionally, mechanistic details of how Neat1 stabilizes Hif1a mRNA— including whether this involves ribonucleoprotein complex formation or paraspeckle-dependent pathways—require further elucidation. Finally, while we demonstrate beneficial effects of Neat1_1 overexpression in aged mice, the translational feasibility, timing, and safety of modulating Neat1_1 in vivo have not yet been tested.

Future research should evaluate cell type–specific roles of Neat1 in the muscle niche, define the molecular interactome of Neat1-Hif1a mRNA complexes, and assess whether targeted delivery of Neat1_1 or Neat1-based therapeutics can enhance recovery in clinically relevant models of peripheral artery disease and age-related sarcopenia. Ultimately, these studies could establish Neat1_1 as a mechanistic target to restore hypoxic resilience and regenerative function in aging muscle.

## Acknowledgement

We gratefully acknowledge the LifeLink Foundation for providing human muscle tissue, which enabled the isolation of human muscle cells. HM and JHY were supported by startup fund from University of Oklahoma.

## Author Contribution

KK was involved in designing the study, conducting experiments, analyzing data, creating figures, and writing the manuscript. TY, HM, DGO, and SL conducted experiments, analyzing data and creating figures. MK, JK, YK, and JL conducted experiments and data analysis. YCJ, JHY, LH, and HJC were involved in designing the study, analyzing data, creating figures, writing the manuscript, and providing funds.

## Notes

### Competing Interest Statement

The authors have declared no competing interest.

## References

1. Mozaffarian, D., et al. Executive Summary: Heart Disease and Stroke Statistics— 2015 Update: A Report From the American Heart Association. Circulation 131, 434–441 (2015).

2. Barnes, J.A., Eid, M.A., Creager, M.A. & Goodney, P.P. Epidemiology and Risk of Amputation in Patients With Diabetes Mellitus and Peripheral Artery Disease. Arteriosclerosis, thrombosis, and vascular biology 40, 1808–1817 (2020).

3. Beckman, J.A., Schneider, P.A. & Conte, M.S. Advances in Revascularization for Peripheral Artery Disease: Revascularization in PAD. Circulation Research 128, 1885–1912 (2021).

4. Defraigne, J.O. & Pincemail, J. Local and systemic consequences of severe ischemia and reperfusion of the skeletal muscle. Physiopathology and prevention. Acta Chir Belg 98, 176–186 (1998).

5. Han, J., Luo, L., Marcelina, O., Kasim, V. & Wu, S. Therapeutic angiogenesis-based strategy for peripheral artery disease. Theranostics 12, 5015–5033 (2022).

6. Ferrari, R. et al. Inflammatory Caspase Activity Mediates HMGB1 Release and Differentiation in Myoblasts Affected by Peripheral Arterial Disease. Cells 11 (2022).

7. McDermott, M.M. Peripheral Artery Disease: Past and Future. Circulation 149, 1151–1153 (2024).

8. Jazwa, A. et al. Limb ischemia and vessel regeneration: Is there a role for VEGF? Vascul Pharmacol 86, 18–30 (2016).

9. Grochot-Przeczek, A., Dulak, J. & Jozkowicz, A. Therapeutic angiogenesis for revascularization in peripheral artery disease. Gene 525, 220–228 (2013).

10. Moroz, E. et al. Real-time imaging of HIF-1alpha stabilization and degradation. PLoS One 4, e5077 (2009).

11. Serocki, M. et al. miRNAs regulate the HIF switch during hypoxia: a novel therapeutic target. Angiogenesis 21, 183–202 (2018).

12. Salceda, S. & Caro, J. Hypoxia-inducible factor 1alpha (HIF-1alpha) protein is rapidly degraded by the ubiquitin-proteasome system under normoxic conditions. Its stabilization by hypoxia depends on redox-induced changes. J Biol Chem 272, 22642–22647 (1997).

13. Huang, L.E., Arany, Z., Livingston, D.M. & Bunn, H.F. Activation of hypoxia-inducible transcription factor depends primarily upon redox-sensitive stabilization of its alpha subunit. J Biol Chem 271, 32253–32259 (1996).

14. Yamamoto, H. et al. Efficacy and Safety of Molidustat for Anemia in ESA-Naive Nondialysis Patients: A Randomized, Phase 3 Trial. Am J Nephrol 52, 871–883 (2021).

15. Eckardt, K.U. et al. Safety and Efficacy of Vadadustat for Anemia in Patients Undergoing Dialysis. N Engl J Med 384, 1601–1612 (2021).

16. Chen, N. et al. Roxadustat for Anemia in Patients with Kidney Disease Not Receiving Dialysis. N Engl J Med 381, 1001–1010 (2019).

17. Singh, A.K. et al. Daprodustat for the Treatment of Anemia in Patients Undergoing Dialysis. N Engl J Med 385, 2325–2335 (2021).

18. Bao, W. et al. Chronic inhibition of hypoxia-inducible factor prolyl 4-hydroxylase improves ventricular performance, remodeling, and vascularity after myocardial infarction in the rat. J Cardiovasc Pharmacol 56, 147–155 (2010).

19. Chen, R.L. et al. Roles of individual prolyl-4-hydroxylase isoforms in the first 24 hours following transient focal cerebral ischaemia: insights from genetically modified mice. J Physiol 590, 4079–4091 (2012).

20. Billin, A.N. et al. HIF prolyl hydroxylase inhibition protects skeletal muscle from eccentric contraction-induced injury. Skelet Muscle 8, 35 (2018).

21. Olson, E. et al. Short-term treatment with a novel HIF-prolyl hydroxylase inhibitor (GSK1278863) failed to improve measures of performance in subjects with claudication-limited peripheral artery disease. Vasc Med 19, 473–482 (2014).

22. Cirillo, F. et al. Human Sarcopenic Myoblasts Can Be Rescued by Pharmacological Reactivation of HIF-1alpha. Int J Mol Sci 23 (2022).

23. Endo, Y. et al. Loss of ARNT in skeletal muscle limits muscle regeneration in aging. FASEB J 34, 16086–16104 (2020).

24. Yang, X., Yang, S., Wang, C. & Kuang, S. The hypoxia-inducible factors HIF1alpha and HIF2alpha are dispensable for embryonic muscle development but essential for postnatal muscle regeneration. J Biol Chem 292, 5981–5991 (2017).

25. Onoguchi-Mizutani, R. & Akimitsu, N. Long noncoding RNA and phase separation in cellular stress response. J Biochem 171, 269–276 (2022).

26. Choudhry, H. et al. Tumor hypoxia induces nuclear paraspeckle formation through HIF-2alpha dependent transcriptional activation of NEAT1 leading to cancer cell survival. Oncogene 34, 4482–4490 (2015).

27. Godet, A.C. et al. Long non-coding RNA Neat1 and paraspeckle components are translational regulators in hypoxia. Elife 11 (2022).

28. Lellahi, S.M. et al. The long noncoding RNA NEAT1 and nuclear paraspeckles are up-regulated by the transcription factor HSF1 in the heat shock response. J Biol Chem 293, 18965–18976 (2018).

29. Fox, A.H., Nakagawa, S., Hirose, T. & Bond, C.S. Paraspeckles: Where Long Noncoding RNA Meets Phase Separation. Trends Biochem Sci 43, 124–135 (2018).

30. Clemson, C.M. et al. An architectural role for a nuclear noncoding RNA: NEAT1 RNA is essential for the structure of paraspeckles. Mol Cell 33, 717–726 (2009).

31. Sasaki, Y.T., Ideue, T., Sano, M., Mituyama, T. & Hirose, T. MENepsilon/beta noncoding RNAs are essential for structural integrity of nuclear paraspeckles. Proc Natl Acad Sci U S A 106, 2525–2530 (2009).

32. Sunwoo, H. et al. MEN epsilon/beta nuclear-retained non-coding RNAs are up-regulated upon muscle differentiation and are essential components of paraspeckles. Genome Res 19, 347–359 (2009).

33. Fox, A.H. & Lamond, A.I. Paraspeckles. Cold Spring Harb Perspect Biol 2, a000687 (2010).

34. Scholda, J., Nguyen, T.T.A. & Kopp, F. Long noncoding RNAs as versatile molecular regulators of cellular stress response and homeostasis. Hum Genet (2023).

35. Morrison, S.J. Neuronal potential and lineage determination by neural stem cells. Curr Opin Cell Biol 13, 666–672 (2001).

36. An, H., Williams, N.G. & Shelkovnikova, T.A. NEAT1 and paraspeckles in neurodegenerative diseases: A missing lnc found? Noncoding RNA Res 3, 243–252 (2018).

37. Lulli, V. et al. Mir-370-3p Impairs Glioblastoma Stem-Like Cell Malignancy Regulating a Complex Interplay between HMGA2/HIF1A and the Oncogenic Long Non-Coding RNA (lncRNA) NEAT1. Int J Mol Sci 21 (2020).

38. Yu, L. et al. Silencing long non-coding RNA NEAT1 suppresses the tumorigenesis of infantile hemangioma by competitively binding miR-33a-5p to stimulate HIF1alpha/NF-kappaB pathway. Mol Med Rep 22, 3358–3366 (2020).

39. Jiang, P. et al. LncRNA NEAT1 contributes to the acquisition of a tumor like-phenotype induced by PM 2.5 in lung bronchial epithelial cells via HIF-1alpha activation. Environ Sci Pollut Res Int 28, 43382–43393 (2021).

40. Park, M.K. et al. NEAT1 is essential for metabolic changes that promote breast cancer growth and metastasis. Cell Metab 33, 2380–2397 e2389 (2021).

41. Liu, Y. et al. The long non-coding RNA NEAT1 promotes the progression of human ovarian cancer through targeting miR-214-3p and regulating angiogenesis. J Ovarian Res 16, 219 (2023).

42. Zhang, Q. et al. Hypoxia-Induced lncRNA-NEAT1 Sustains the Growth of Hepatocellular Carcinoma via Regulation of miR-199a-3p/UCK2. Front Oncol 10, 998 (2020).

43. Zhong, J. et al. The long non-coding RNA Neat1 is an important mediator of the therapeutic effect of bexarotene on traumatic brain injury in mice. Brain Behav Immun 65, 183–194 (2017).

44. Nakagawa, S. et al. The lncRNA Neat1 is required for corpus luteum formation and the establishment of pregnancy in a subpopulation of mice. Development 141, 4618–4627 (2014).

45. Foltz, S.J., Hartzell, C.H. & Choo, H.J. In vivo Electroporation of Skeletal Muscle Fibers in Mice. Bio Protoc 13, e4759 (2023).

46. Mohiuddin, M. et al. Critical Limb Ischemia Induces Remodeling of Skeletal Muscle Motor Unit, Myonuclear-, and Mitochondrial-Domains. Sci Rep 9, 9551 (2019).

47. Kim, E. et al. Fibroadipogenic Progenitors Regulate the Basal Proliferation of Satellite Cells and Homeostasis of Pharyngeal Muscles via HGF Secretion. Front Cell Dev Biol 10, 875209 (2022).

48. Helisch, A. et al. Impact of mouse strain differences in innate hindlimb collateral vasculature. Arterioscler Thromb Vasc Biol 26, 520–526 (2006).

49. Keith, B., Johnson, R.S. & Simon, M.C. HIF1alpha and HIF2alpha: sibling rivalry in hypoxic tumour growth and progression. Nat Rev Cancer 12, 9–22 (2011).

50. Higashimura, Y. et al. Up-regulation of glyceraldehyde-3-phosphate dehydrogenase gene expression by HIF-1 activity depending on Sp1 in hypoxic breast cancer cells. Arch Biochem Biophys 509, 1–8 (2011).

51. Semenza, G.L. Hypoxia, clonal selection, and the role of HIF-1 in tumor progression. Crit Rev Biochem Mol Biol 35, 71–103 (2000).

52. Bosch-Marce, M. et al. Effects of aging and hypoxia-inducible factor-1 activity on angiogenic cell mobilization and recovery of perfusion after limb ischemia. Circ Res 101, 1310–1318 (2007).

53. Loinard, C. et al. Inhibition of prolyl hydroxylase domain proteins promotes therapeutic revascularization. Circulation 120, 50–59 (2009).

